## Supplementary material for "Adventitial fibroblasts direct smooth muscle cell-state transition in pulmonary vascular disease": Supplemantary Figures and Tables

**This PDF file includes:**

Figures S1 to S8

Tables S1 to S3

**Other Supplementary Materials for this manuscript include the following:**

Data S1 to S4

A

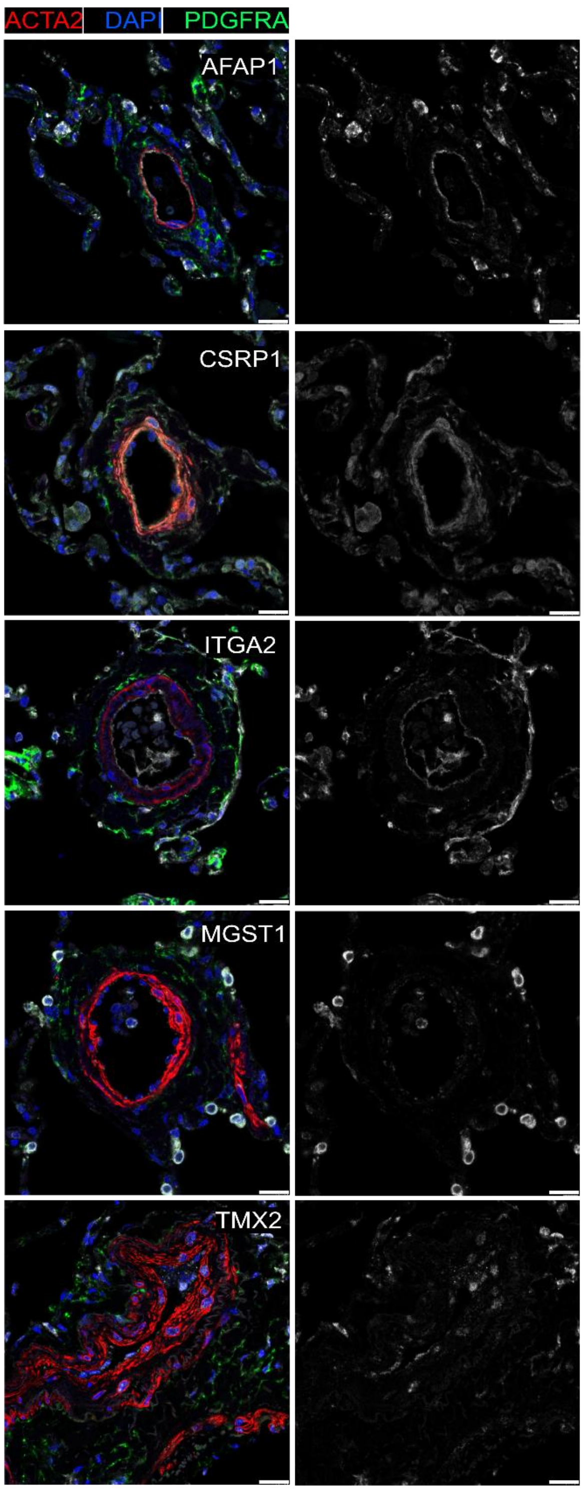


B

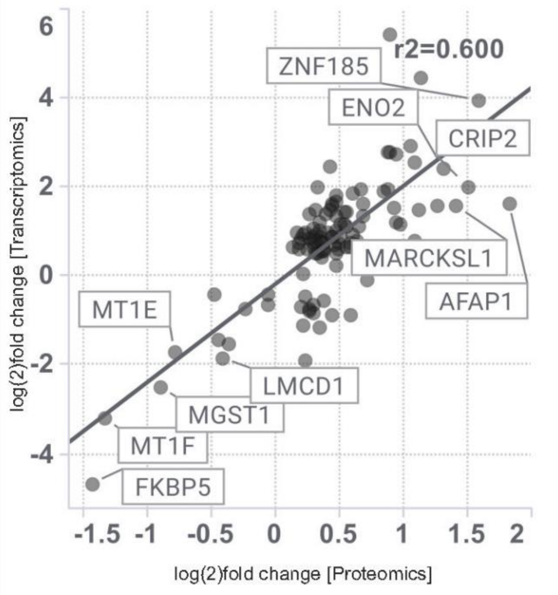


| | proteomics | | transcriptomics | |
|---|---|---|---|---|
| | LFC | -log p | LFC | -log p |
| AFAP1 | 1,82 | 3,44 | 1,63 | 2,27 |
| CSRP1 | 1,25 | 3,11 | 1,58 | 1,86 |
| ITGA2 | 1,12 | 5,73 | 4,49 | 5,68 |
| MGST1 | -0,9 | 2,75 | -2,51 | 6,63 |
| TMX2 | -0,68 | 2,71 | 0,08 | 0,11 |

C

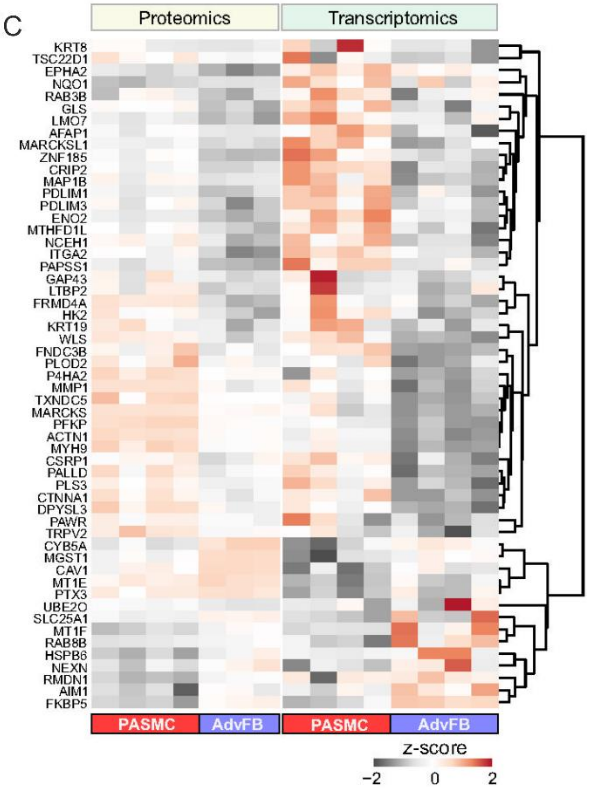

**Figure S1. Validation and correlation between transcriptomics and proteomics.**

A) Representative immunofluorescent images of PASMC-enriched (Actin Filament Associated Protein 1 – AFAP1, Cysteine and Glycine Rich Protein 1 – CSRP1, Integrin Alpha 2 - ITGA2) and PAAF-enriched (Microsomal Glutathione S-Transferase 1 – MGST1, Thioredoxin Related Transmembrane Protein 2 – TMX2) molecules on human donor pulmonary arteries. Co-staining with ACTA2 as PASMC marker and PDGFRA as fibroblast marker. DAPI as nuclear counterstain. White bar depicting 20 μm. B) Correlation plot displaying the linear relationship between the $log_2$ fold change observed in transcriptomics and proteomics. Plotted are all genes that were significantly regulated between PASMC and PAAF in both -omics approaches (-log10(P)>2 in the pair-wise comparison between donor PASMC and PAAF) and genes where significance was given by ANOVA (-log10(P)>2 when comparing all four groups. The linear Pearson correlation is marked by the black line and the corresponding $R^2$. C) Heatmap showing the single expression values for the genes that were found to be significantly regulated between PASMC and PAAF in either transcriptomics or proteomics.

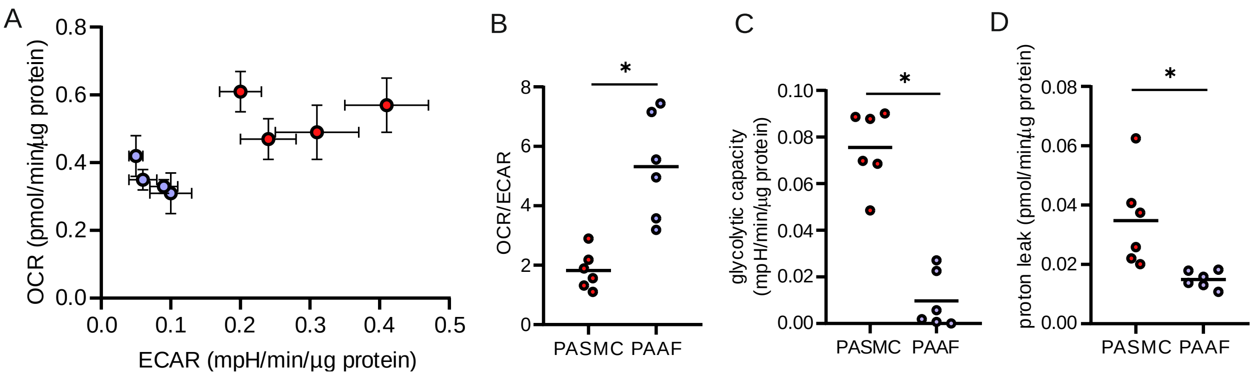


**Figure S2. Real-time metabolic analysis between donor PASMC and PAAF.**

A) Representative energy map of oxygen consumption and extracellular acidification rate. B) Ratio of oxygen consumption rate (OCR) to extracellular acidification rate (ECAR). C) Measured glycolytic capacity (mpH/min) normalized to total protein content. D) Calculated mitochondrial proton leak (pmol/min) normalized to total protein content. Mann-Whitney test, $p<0.05$.

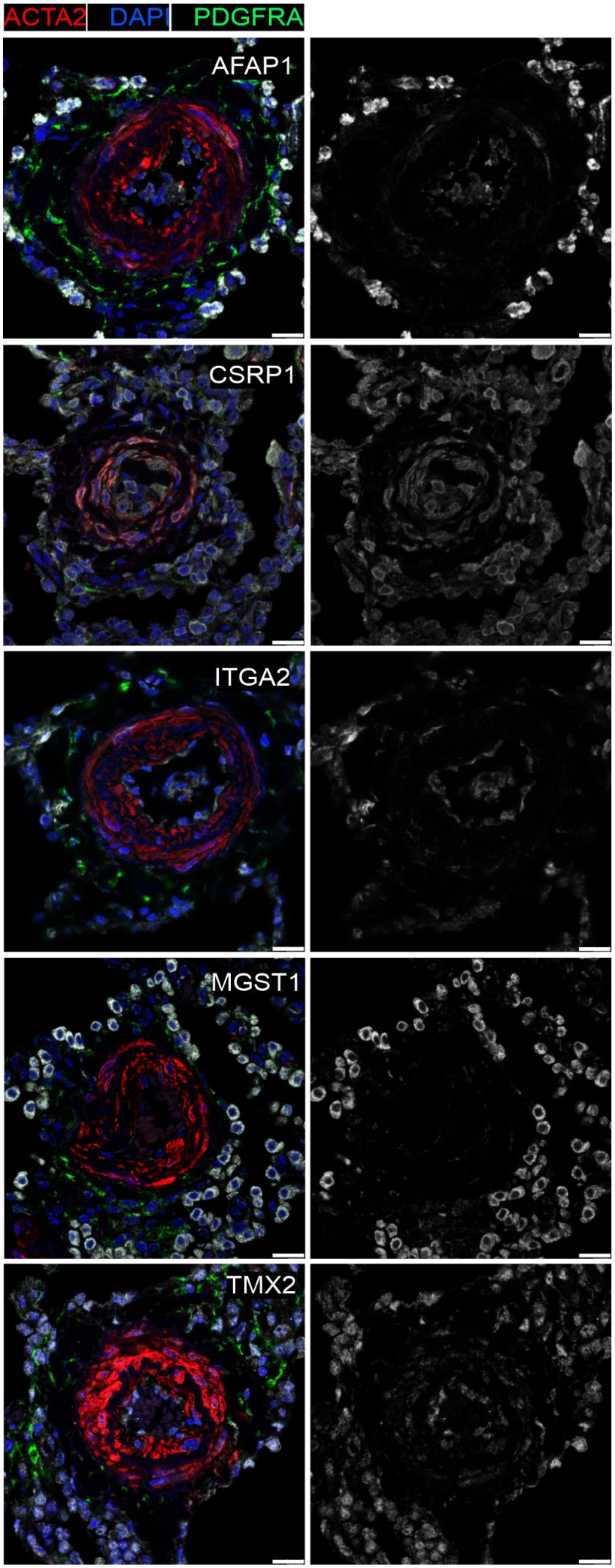
ACTA2
DAPI
PDGFRA
AFAP1
CSRP1
ITGA2
MGST1
TMX2

**Figure S3. Validation of omics markers in IPAH.**

Representative immunofluorescent images of PASMC-enriched (AFAP1, CSRP1, ITGA2) and PAAF-enriched (MGST1, TMX2,) molecules on human IPAH pulmonary arteries. Co-staining with ACTA2 as PASMC marker and PDGFRA as fibroblast marker. DAPI as nuclear counterstain. White bar depicting 20 μm.

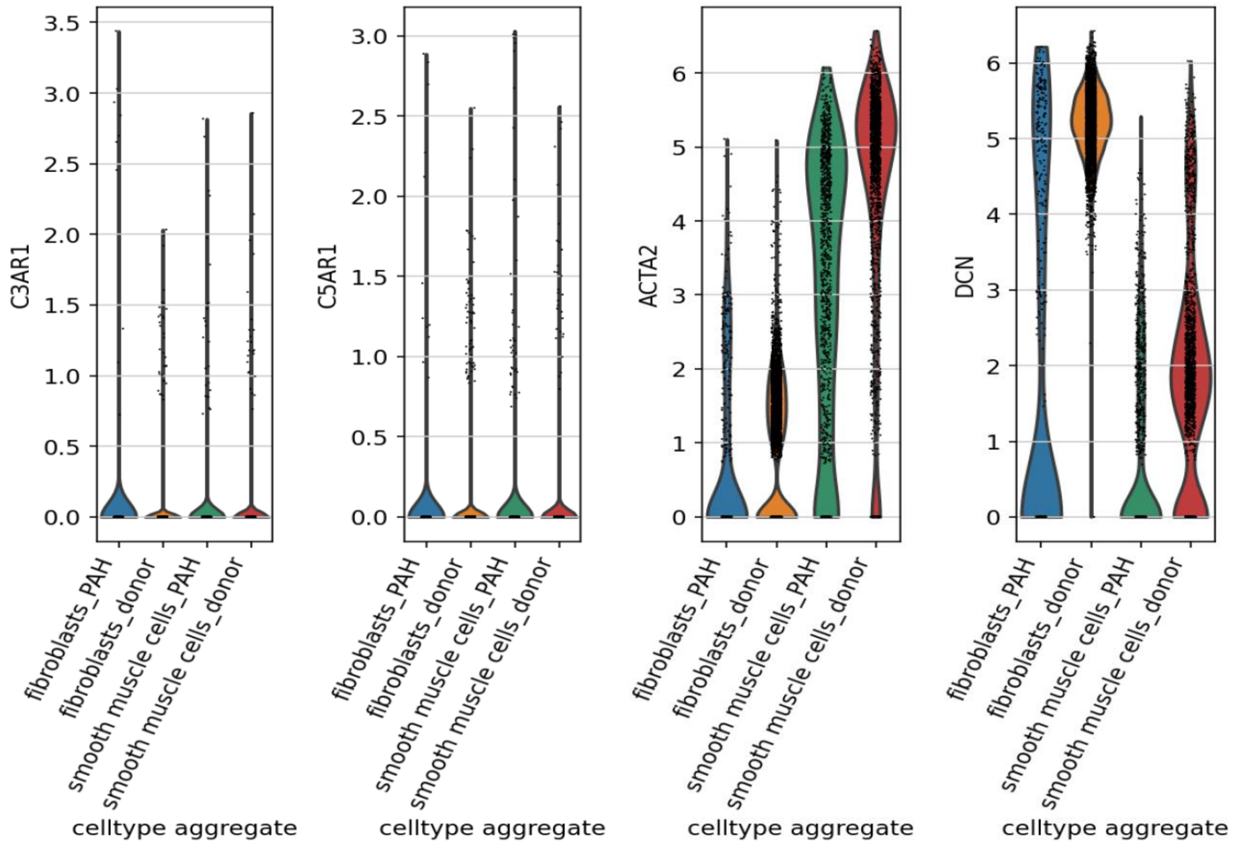


**Figure S4. PASMC and PAAF expression of anaphylatoxin receptors.**

Violin plots showing the expression of complement component 3a receptor 1 (C3AR1) and complement component 5a receptor 1 (C5AR1) in adventitial fibroblasts and smooth muscle cells clusters, identified by DCN and ACTA2 canonical marker expression, from human pulmonary arteries single cell RNA-Seq dataset (GSE210248).

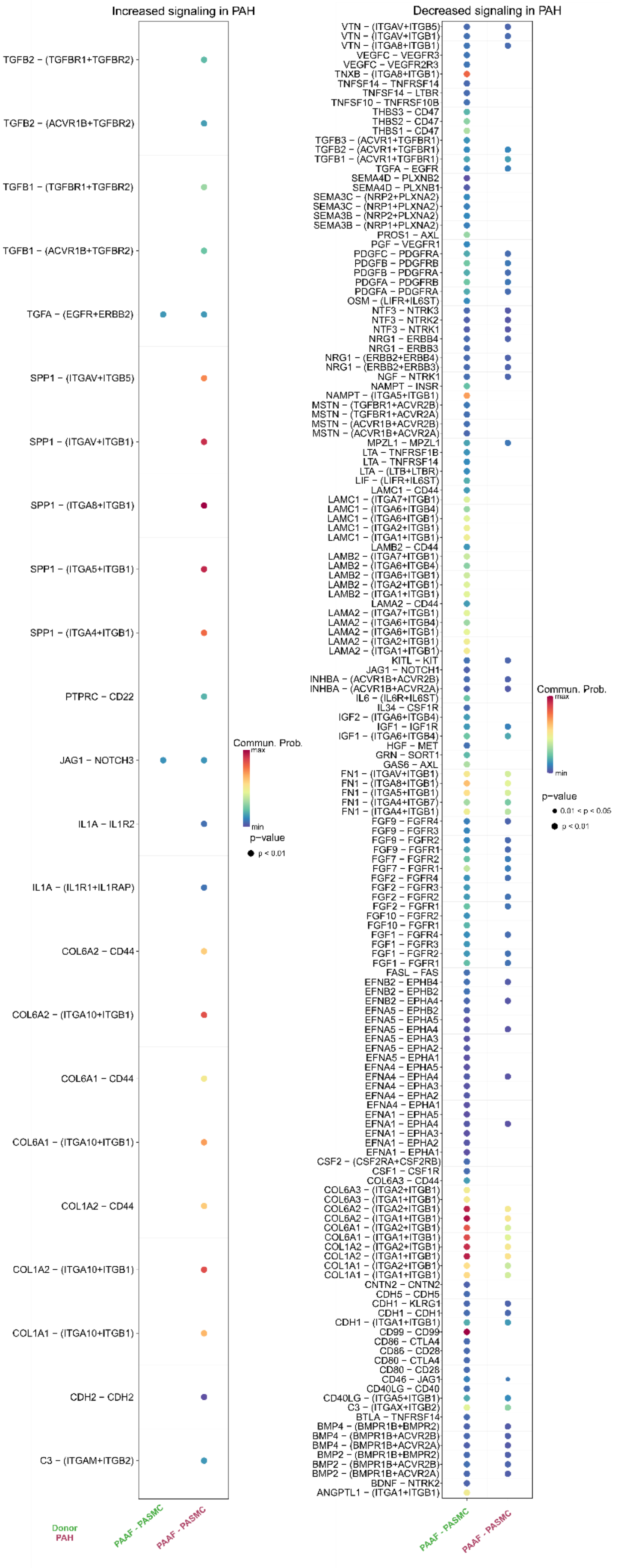

Increased signaling in PAH
TGFB2 − (TGFBR1+TGFBR2)
TGFB2 − (ACVR1B+TGFBR2)
TGFB1 − (TGFBR1+TGFBR2)
TGFB1 − (ACVR1B+TGFBR2)
TGFA − (EGFR+ERBB2)
SPP1 − (ITGAV+ITGB5)
SPP1 − (ITGAV+ITGB1)
SPP1 − (ITGA8+ITGB1)
SPP1 − (ITGA5+ITGB1)
SPP1 − (ITGA4+ITGB1)
PTPRC − CD22
JAG1 − NOTCH3
IL1A − IL1R2
IL1A − (IL1R1+IL1RAP)
COL6A2 − CD44
COL6A2 − (ITGA10+ITGB1)
COL6A1 − CD44
COL6A1 − (ITGA10+ITGB1)
COL1A2 − CD44
COL1A2 − (ITGA10+ITGB1)
COL1A1 − (ITGA10+ITGB1)
CDH2 − CDH2
C3 − (ITGAM+ITGB2)
Commun. Prob.
max
min
p−value
p < 0.01
Donor
PAH
PAAF - PASMC
PAAF - PASMC
Decreased signaling in PAH
VTN − (ITGAV+ITGB5)
VTN − (ITGAV+ITGB1)
VTN − (ITGA8+ITGB1)
VEGFC − VEGFR3
VEGFC − VEGFR2R3
TNXB − (ITGA8+ITGB1)
TNFSF14 − TNFRSF14
TNFSF14 − LTBR
TNFSF10 − TNFRSF10B
THBS3 − CD47
THBS2 − CD47
THBS1 − CD47
TGFB3 − (ACVR1+TGFBR1)
TGFB2 − (ACVR1+TGFBR1)
TGFB1 − (ACVR1+TGFBR1)
TGFA − EGFR
SEMA4D − PLXNB2
SEMA4D − PLXNB1
SEMA3C − (NRP2+PLXNA2)
SEMA3C − (NRP1+PLXNA2)
SEMA3B − (NRP2+PLXNA2)
SEMA3B − (NRP1+PLXNA2)
PROS1 − AXL
PGF − VEGFR1
PDGFC − PDGFRA
PDGFB − PDGFRB
PDGFB − PDGFRA
PDGFA − PDGFRB
PDGFA − PDGFRA
OSM − (LIFR+IL6ST)
NTF3 − NTRK3
NTF3 − NTRK2
NTF3 − NTRK1
NRG1 − ERBB4
NRG1 − ERBB3
NRG1 − (ERBB2+ERBB4)
NRG1 − (ERBB2+ERBB3)
NGF − NTRK1
NAMPT − INSR
NAMPT − (ITGA5+ITGB1)
MSTN − (TGFBR1+ACVR2B)
MSTN − (TGFBR1+ACVR2A)
MSTN − (ACVR1B+ACVR2B)
MSTN − (ACVR1B+ACVR2A)
MPZL1 − MPZL1
LTA − TNFRSF1B
LTA − TNFRSF14
LTA − (LTB+LTBR)
LIF − (LIFR+IL6ST)
LAMC1 − CD44
LAMC1 − (ITGA7+ITGB1)
LAMC1 − (ITGA6+ITGB4)
LAMC1 − (ITGA6+ITGB1)
LAMC1 − (ITGA2+ITGB1)
LAMC1 − (ITGA1+ITGB1)
LAMB2 − CD44
LAMB2 − (ITGA7+ITGB1)
LAMB2 − (ITGA6+ITGB4)
LAMB2 − (ITGA6+ITGB1)
LAMB2 − (ITGA2+ITGB1)
LAMB2 − (ITGA1+ITGB1)
LAMA2 − CD44
LAMA2 − (ITGA7+ITGB1)
LAMA2 − (ITGA6+ITGB4)
LAMA2 − (ITGA6+ITGB1)
LAMA2 − (ITGA2+ITGB1)
LAMA2 − (ITGA1+ITGB1)
KITL − KIT
JAG1 − NOTCH1
INHBA − (ACVR1B+ACVR2B)
INHBA − (ACVR1B+ACVR2A)
IL6 − (IL6R+IL6ST)
IL34 − CSF1R
IGF2 − (ITGA6+ITGB4)
IGF1 − IGF1R
IGF1 − (ITGA6+ITGB4)
HGF − MET
GRN − SORT1
GAS6 − AXL
FN1 − (ITGAV+ITGB1)
FN1 − (ITGA8+ITGB1)
FN1 − (ITGA5+ITGB1)
FN1 − (ITGA4+ITGB7)
FN1 − (ITGA4+ITGB1)
FGF9 − FGFR4
FGF9 − FGFR3
FGF9 − FGFR2
FGF9 − FGFR1
FGF7 − FGFR2
FGF7 − FGFR1
FGF2 − FGFR4
FGF2 − FGFR3
FGF2 − FGFR2
FGF2 − FGFR1
FGF10 − FGFR2
FGF10 − FGFR1
FGF1 − FGFR4
FGF1 − FGFR3
FGF1 − FGFR2
FGF1 − FGFR1
FASL − FAS
EFNB2 − EPHB4
EFNB2 − EPHB2
EFNB2 − EPHA4
EFNA5 − EPHB2
EFNA5 − EPHA5
EFNA5 − EPHA4
EFNA5 − EPHA3
EFNA5 − EPHA2
EFNA5 − EPHA1
EFNA4 − EPHA5
EFNA4 − EPHA4
EFNA4 − EPHA3
EFNA4 − EPHA2
EFNA4 − EPHA1
EFNA1 − EPHA5
EFNA1 − EPHA4
EFNA1 − EPHA3
EFNA1 − EPHA2
EFNA1 − EPHA1
CSF2 − (CSF2RA+CSF2RB)
CSF1 − CSF1R
COL6A3 − CD44
COL6A3 − (ITGA2+ITGB1)
COL6A3 − (ITGA1+ITGB1)
COL6A2 − (ITGA2+ITGB1)
COL6A2 − (ITGA1+ITGB1)
COL6A1 − (ITGA2+ITGB1)
COL6A1 − (ITGA1+ITGB1)
COL1A2 − (ITGA2+ITGB1)
COL1A2 − (ITGA1+ITGB1)
COL1A1 − (ITGA2+ITGB1)
COL1A1 − (ITGA1+ITGB1)
CNTN2 − CNTN2
CDH5 − CDH5
CDH1 − KLRG1
CDH1 − CDH1
CDH1 − (ITGA1+ITGB1)
CD99 − CD99
CD86 − CTLA4
CD86 − CD28
CD80 − CTLA4
CD80 − CD28
CD46 − JAG1
CD40LG − CD40
CD40LG − (ITGA5+ITGB1)
C3 − (ITGAX+ITGB2)
BTLA − TNFRSF14
BMP4 − (BMPR1B+BMPR2)
BMP4 − (BMPR1B+ACVR2B)
BMP4 − (BMPR1B+ACVR2A)
BMP2 − (BMPR1B+BMPR2)
BMP2 − (BMPR1B+ACVR2B)
BMP2 − (BMPR1B+ACVR2A)
BDNF − NTRK2
ANGPTL1 − (ITGA1+ITGB1)
Commun. Prob.
max
min
p−value
0.01 < p < 0.05
p < 0.01
PAAF - PASMC
PAAF - PASMC

**Figure S5. Differential cell-cell communication analysis with PAAF as sender cell type.**

A comparison of cell-cell-communication in donor and PAH between fibroblasts and smooth muscle cells was performed based on the results from the CellChat analysis. PAAF were set as sender cell type whereas PASMC were set as receiver cell type. The two columns on the left side depict the increased signaling interactions and their corresponding ligand-receptor pairs (L-R-pairs), the two columns on the right showing decreased signaling and the corresponding L-R-pairs. Dot size depicting significance and color representing communication probability.

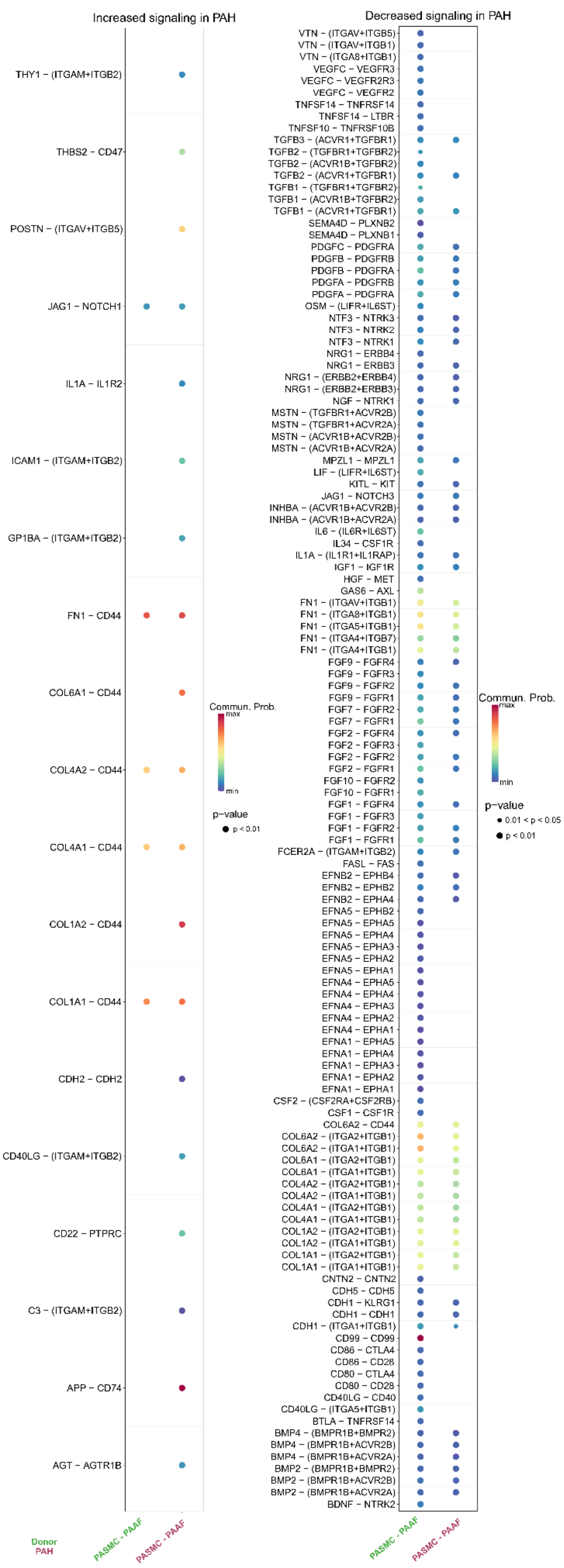
Increased signaling in PAH
THY1 – (ITGAM+ITGB2)
THBS2 – CD47
POSTN – (ITGAV+ITGB5)
JAG1 – NOTCH1
IL1A – IL1R2
ICAM1 – (ITGAM+ITGB2)
GP1BA – (ITGAM+ITGB2)
FN1 – CD44
COL6A1 – CD44
COL4A2 – CD44
COL4A1 – CD44
COL1A2 – CD44
COL1A1 – CD44
CDH2 – CDH2
CD40LG – (ITGAM+ITGB2)
CD22 – PTPRC
C3 – (ITGAM+ITGB2)
APP – CD74
AGT – AGTR1B
Commun. Prob.
max
min
p−value
p < 0.01
Donor
PAH
PASMC - PAAF
PASMC - PAAF
Decreased signaling in PAH
VTN – (ITGAV+ITGB5)
VTN – (ITGAV+ITGB1)
VTN – (ITGA8+ITGB1)
VEGFC – VEGFR3
VEGFC – VEGFR2R3
VEGFC – VEGFR2
TNFSF14 – TNFRSF14
TNFSF14 – LTBR
TNFSF10 – TNFRSF10B
TGFB3 – (ACVR1+TGFBR1)
TGFB2 – (TGFBR1+TGFBR2)
TGFB2 – (ACVR1B+TGFBR2)
TGFB2 – (ACVR1+TGFBR1)
TGFB1 – (TGFBR1+TGFBR2)
TGFB1 – (ACVR1B+TGFBR2)
TGFB1 – (ACVR1+TGFBR1)
SEMA4D – PLXNB2
SEMA4D – PLXNB1
PDGFC – PDGFRA
PDGFB – PDGFRB
PDGFB – PDGFRA
PDGFA – PDGFRB
PDGFA – PDGFRA
OSM – (LIFR+IL6ST)
NTF3 – NTRK3
NTF3 – NTRK2
NTF3 – NTRK1
NRG1 – ERBB4
NRG1 – ERBB3
NRG1 – (ERBB2+ERBB4)
NRG1 – (ERBB2+ERBB3)
NGF – NTRK1
MSTN – (TGFBR1+ACVR2B)
MSTN – (TGFBR1+ACVR2A)
MSTN – (ACVR1B+ACVR2B)
MSTN – (ACVR1B+ACVR2A)
MPZL1 – MPZL1
LIF – (LIFR+IL6ST)
KITL – KIT
JAG1 – NOTCH3
INHBA – (ACVR1B+ACVR2B)
INHBA – (ACVR1B+ACVR2A)
IL6 – (IL6R+IL6ST)
IL34 – CSF1R
IL1A – (IL1R1+IL1RAP)
IGF1 – IGF1R
HGF – MET
GAS6 – AXL
FN1 – (ITGAV+ITGB1)
FN1 – (ITGA8+ITGB1)
FN1 – (ITGA5+ITGB1)
FN1 – (ITGA4+ITGB7)
FN1 – (ITGA4+ITGB1)
FGF9 – FGFR4
FGF9 – FGFR3
FGF9 – FGFR2
FGF9 – FGFR1
FGF7 – FGFR2
FGF7 – FGFR1
FGF2 – FGFR4
FGF2 – FGFR3
FGF2 – FGFR2
FGF2 – FGFR1
FGF10 – FGFR2
FGF10 – FGFR1
FGF1 – FGFR4
FGF1 – FGFR3
FGF1 – FGFR2
FGF1 – FGFR1
FCER2A – (ITGAM+ITGB2)
FASL – FAS
EFNB2 – EPHB4
EFNB2 – EPHB2
EFNB2 – EPHA4
EFNA5 – EPHB2
EFNA5 – EPHA5
EFNA5 – EPHA4
EFNA5 – EPHA3
EFNA5 – EPHA2
EFNA5 – EPHA1
EFNA4 – EPHA5
EFNA4 – EPHA4
EFNA4 – EPHA3
EFNA4 – EPHA2
EFNA4 – EPHA1
EFNA1 – EPHA5
EFNA1 – EPHA4
EFNA1 – EPHA3
EFNA1 – EPHA2
EFNA1 – EPHA1
CSF2 – (CSF2RA+CSF2RB)
CSF1 – CSF1R
COL6A2 – CD44
COL6A2 – (ITGA2+ITGB1)
COL6A2 – (ITGA1+ITGB1)
COL6A1 – (ITGA2+ITGB1)
COL6A1 – (ITGA1+ITGB1)
COL4A2 – (ITGA2+ITGB1)
COL4A2 – (ITGA1+ITGB1)
COL4A1 – (ITGA2+ITGB1)
COL4A1 – (ITGA1+ITGB1)
COL1A2 – (ITGA2+ITGB1)
COL1A2 – (ITGA1+ITGB1)
COL1A1 – (ITGA2+ITGB1)
COL1A1 – (ITGA1+ITGB1)
CNTN2 – CNTN2
CDH5 – CDH5
CDH1 – KLRG1
CDH1 – CDH1
CDH1 – (ITGA1+ITGB1)
CD99 – CD99
CD86 – CTLA4
CD86 – CD28
CD80 – CTLA4
CD80 – CD28
CD40LG – CD40
CD40LG – (ITGA5+ITGB1)
BTLA – TNFRSF14
BMP4 – (BMPR1B+BMPR2)
BMP4 – (BMPR1B+ACVR2B)
BMP4 – (BMPR1B+ACVR2A)
BMP2 – (BMPR1B+BMPR2)
BMP2 – (BMPR1B+ACVR2B)
BMP2 – (BMPR1B+ACVR2A)
BDNF – NTRK2
Commun. Prob.
max
min
p−value
0.01 < p < 0.05
p < 0.01
PASMC - PAAF
PASMC - PAAF

**Figure S6. Differential cell-cell communication analysis with PASMC as sender cell type.**

A comparison of cell-cell-communication in donor and PAH between fibroblasts and smooth muscle cells was performed based on the results from the CellChat analysis. PAAF were set as sender cell type whereas PASMC were set as receiver cell type. The two columns on the left side depict the increased signaling interactions and their corresponding ligand-receptor pairs (L-R-pairs), the two columns on the right showing decreased signaling and the corresponding L-R-pairs. Dot size depicting significance and color representing communication probability.

A

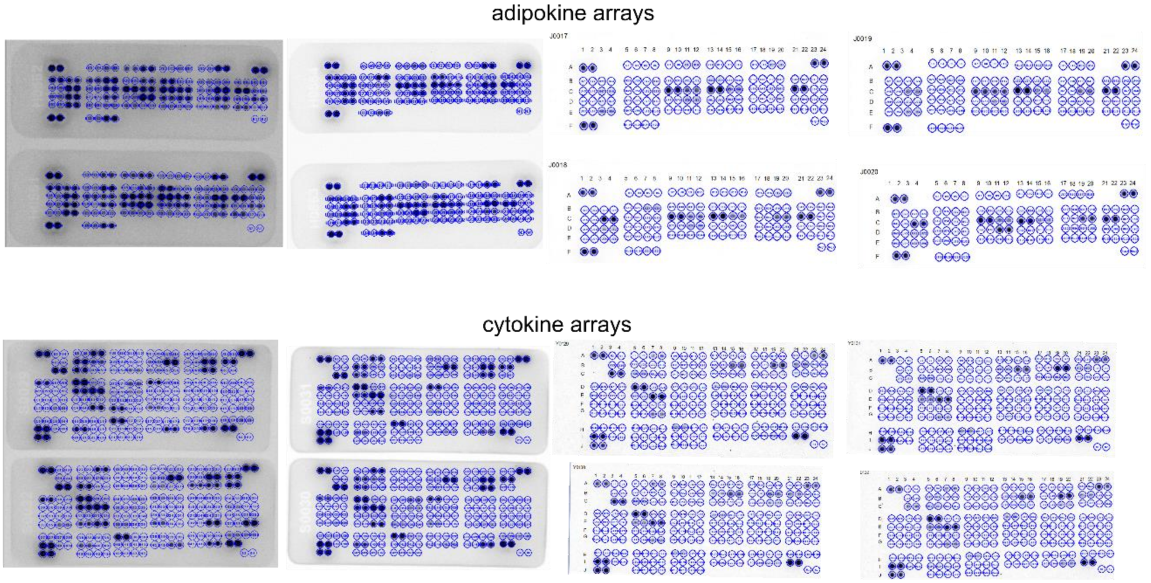

adipokine arrays
cytokine arrays


B

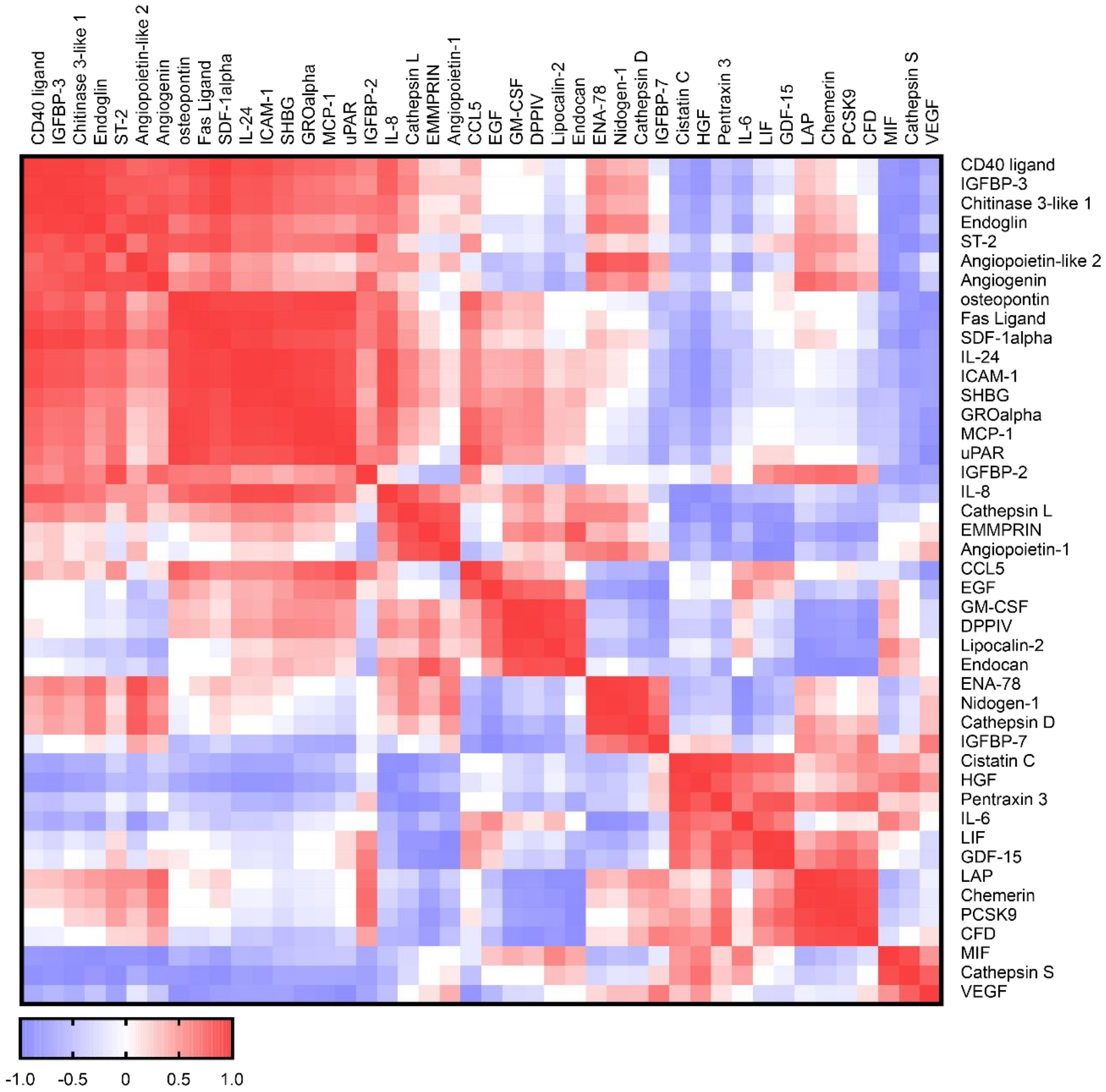

CD40 ligand
IGFBP-3
Chitinase 3-like 1
Endoglin
ST-2
Angiopoietin-like 2
Angiogenin
osteopontin
Fas Ligand
SDF-1alpha
IL-24
ICAM-1
SHBG
GROalpha
MCP-1
uPAR
IGFBP-2
IL-8
Cathepsin L
EMMPRIN
Angiopoietin-1
CCL5
EGF
GM-CSF
DPPIV
Lipocalin-2
Endocan
ENA-78
Nidogen-1
Cathepsin D
IGFBP-7
Cistatin C
HGF
Pentraxin 3
IL-6
LIF
GDF-15
LAP
Chemerin
PCSK9
CFD
MIF
Cathepsin S
VEGF
-1.0
-0.5
0
0.5
1.0

**Figure S7. Similarity matrix of soluble factors measured from PAAF cell culture supernatants.**

A) Proteome Profiler Human XL Cytokine Array Kit (R&D Systems) and Proteome Profiler Human Adipokine Array Kit (R&D Sytems) were used to measure factors secreted by donor and IPAH PAAF. B) Heatmap depicts correlation matrix for soluble factors with pronounced normalized relative expression change between IPAH and donor state. Hierarchically clustering was performed in Clustergrammer.

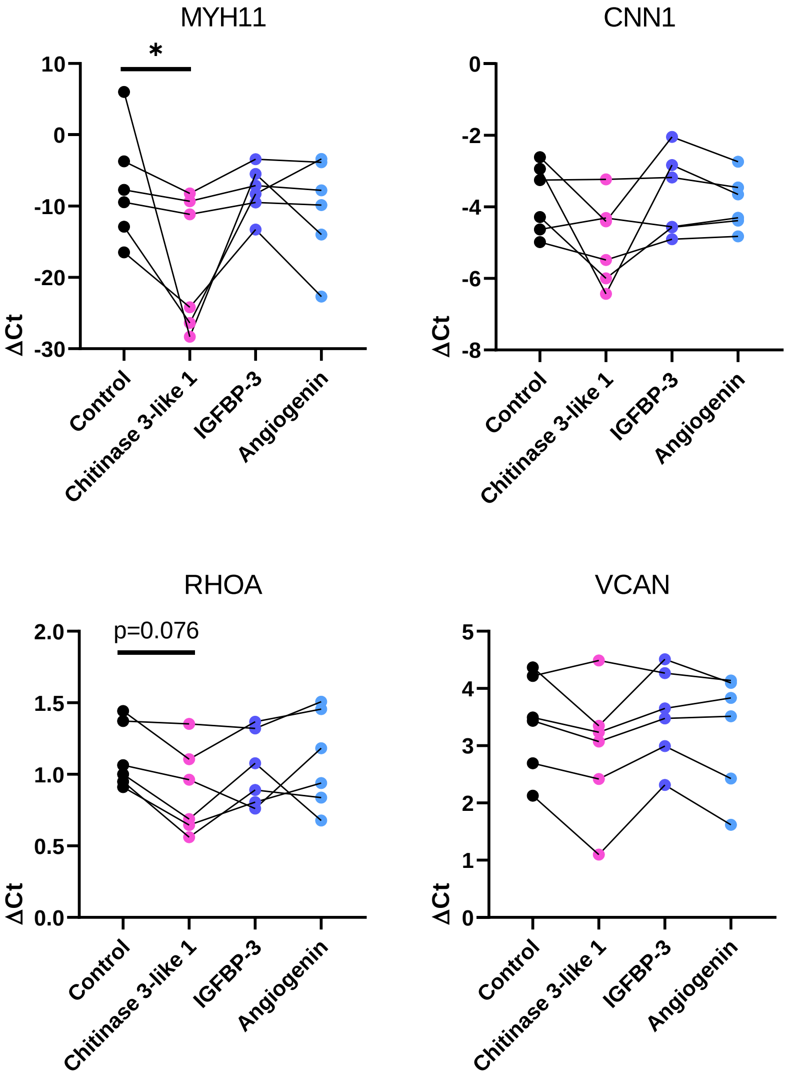


**Figure S8. Gene expression profile of PASMC cell-state markers upon stimulation with additional PAAF ligands.**

Gene expression changes of cell state markers in donor PASMC (n=6) treated 24h with Chitinase 3-like 1 (300 ng/mL), Insulin-like growth factor binding protein-3 (IGFBP-3, 300 ng/mL), and Angiogenin (300 ng/mL). Calponin: CNN1, Ras Homolog Family Member A: RHOA, Versican: VCAN, Smooth muscle myosin heavy chain: MHY11. Friedman test followed by Dunn's multiple comparisons test (*p<0.05).

| Sample | Sex | Age at transplant | Application | mPAP (mmHg) | Cardiac output (L/min) | PAH therapy |
|---|---|---|---|---|---|---|
| donor 1 | male | 45 | omics, cellular assays, GAG quantification | | | |
| donor 2 | female | 58 | omics, cellular assays, GAG quantification | | | |
| donor 3 | female | 44 | omics, GAG quantification, cellular assays | | | |
| donor 4 | male | 23 | omics, GAG quantification, cellular assays | | | |
| donor 5 | female | 24 | GAG quantification, cellular assays | | | |
| donor 6 | female | 40 | GAG quantification, cellular assays | | | |
| donor 7 | male | 62 | GAG quantification, stainings, cellular assays | | | |
| donor 8 | male | 53 | GAG quantification, stainings, cellular assays | | | |
| donor 9 | female | 56 | GAG quantification, stainings, cellular assays | | | |
| donor 10 | female | 76 | GAG quantification, stainings, cellular assays | | | |
| IPAH 1 | female | 32 | omics, GAG quantification, cellular assays | 50 | n.a. | bosentan, sildenafil, epoprostenol |
| IPAH 2 | female | 38 | omics, cellular assays, GAG quantification | 76 | 1.9 | bosentan, sildenafil, treprostinil |
| IPAH 3 | male | 36 | omics, cellular assays, GAG quantification | 62 | 4.3 | macitentan, sildenafil, treprostinil |
| IPAH 4 | male | 42 | omics, GAG quantification, cellular assays | 101 | 3.39 | bosentan, sildenafil, treprostinil |

| | | | | | | |
|---|---|---|---|---|---|---|
| IPAH 5 | female | 27 | GAG quantification, cellular assays | 65 | 3.7 | sildenafil, treprostinil |
| IPAH 6 | female | 52 | GAG quantification, cellular assays | 63 | 2.8 | bosentan, treprostinil |
| IPAH 7 | male | 21 | GAG quantification, stainings, cellular assays | 90 | 3.1 | bosentan, sildenafil |
| IPAH 8 | male | 37 | GAG quantification, stainings, cellular assays | 68 | 5.4 | macitentan, riociguat, treprostinil |
| IPAH 9 | female | 52 | GAG quantification, stainings, cellular assays | 77 | n.a. | bosentan, sildenafil, treprostinil |
| IPAH 10 | female | 32 | GAG quantification, stainings, cellular assays | 74 | n.a. | macitentan, riociguat, treprostinil |

**Table S1. Patient characteristics.**

Age and sex of healthy controls (donors) and patients with pulmonary vascular disease (IPAH) with corresponding clinical data (mean pulmonary arterial pressure, mPAP, cardiac output) and PAH therapy.

| Antibody | Company | Catalog number | Dilution | Detection | ID |
|---|---|---|---|---|---|
| ACTA2-Cy3 | Sigma | C6198 | 1:100 | primary labeled Ab | AB_476856 |
| ACTA2-FITC | Sigma | F3777 | 1:100 | primary labeled Ab | AB_476977 |
| ADH1A/ADH1C | Atlas Antibodies | HPA047814 | 1:300 | Tyramide signal amplification | AB_2680163 |
| AFAP1 | Atlas Antibodies | HPA015642 | 1:200 | Tyramide signal amplification | AB_1844632 |
| CD45 | Abcam | ab10558 | 1:300 | Tyramide signal amplification | AB_442810 |
| CSRP1 | Atlas Antibodies | HPA045617 | 1:100 | Tyramide signal amplification | AB_2679391 |
| ITGA2 | Atlas Antibodies | HPA063556 | 1:100 | Tyramide signal amplification | AB_2685040 |
| MGST1 | ThermoFisher Scientific | PA5-60845 | 1:100 | Tyramide signal amplification | AB_2643943 |
| PCNA | Santa Cruz | sc-7907 | 1:100 | secondary labeled Ab | AB_2160375 |
| PDGFRalpha | Cell Signaling Technologies | 3174 | 1:50 | Tyramide signal amplification | AB_2162345 |
| PDGFRalpha | Abcam | ab203491 | 1:500 | Tyramide signal amplification | AB_2892065 |
| RGS5 | Santa Cruz | sc-514184 | 1:2000 | Tyramide signal amplification | - |
| TMX2 | Atlas Antibodies | HPA063763 | 1:100 | Tyramide signal amplification | AB_2685116 |

| Secondary detection reagents | Company | Catalog number | Dilution |
|---|---|---|---|
| Donkey anti-Rabbit IgG AF-555 | Thermo Fisher Scientific | A-31572 | 1:400 |
| Immpress horse anti- mouse HRP-polymer | Vector Labs | MP-7402 | RTU |
| Immpress horse anti- rabbit HRP-polymer | Vector Labs | MP-7401 | RTU |
| CF405L tyramide dye | Biotium | 92198 | 1:200 |
| CF620R tyramide dye | Biotium | 92194 | 1:200 |
| Opal 7-Color Manual IHC Kit | Akoya Biosciences | SKU NEL811001KT | RTU |
| CF594 TUNEL assay kit | Biotium | 30064 | - |
| DAPI | Thermo Fisher Scientific | 62248 | 1:1000 |

**Table S2. List of antibodies and detection reagents.**

| species | gene symbol | | primer sequence (5´ to 3´) |
|---|---|---|---|
| human | MYH11 | forward | TTGGCTCCCACGATGTAACC |
| | | reverse | AAGCAGCTTCTACAAGCAAAC |
| human | RHOA | forward | AGCAAGCATGTCTTTCCACA |
| | | reverse | GAAGAGGCTGGACTCGGATT |
| human | BGN | forward | CTGGCATCCCCAAAGACCTC |
| | | reverse | CCAGTTCGATGGCCTGGATT |
| human | VCAN | forward | TGTTAATCGTGTGGGCCATGA |
| | | reverse | AGAAGCTGTCTGGCTGGTTG |
| human | CNN1 | forward | CCCCACGACATTTTTGAGGC |
| | | reverse | CACTCCCACGTTCACCTTGT |
| human | PPARG | forward | AGAGCCTTCCAACTCCCTCA |
| | | reverse | TCCGGAAGAAACCCTTGCAT |
| human | TNFRSF11B | forward | CAGTGTCTTTGGTCTCCTGC |
| | | reverse | TCCTCACACAGGGTAACATCTATT |
| human | CCND1 | forward | AGTGGAAACCATCCGCCG |
| | | reverse | TCTGTTCCTCGCAGACCTCCA |
| human | DCN | forward | GCATTCCTCAAGGTCTTCCTCC |
| | | reverse | AGCCATTGTCAACAGCAGAG |
| human | B2M | forward | CCTGGAGGCTATCCAGCGTACTCC |
| | | reverse | TGTCGGATGGATGAAACCCAGACA |
| human | HMBS | forward | CTGCAACGGCGGAAGAAAA |
| | | reverse | AATCTTGTCCCCTGTGGTGG |

**Table S3. List of used primers.**

**Data S1. (separate file)**

List of differentially expressed genes.

**Data S2. (separate file)**

List of detected proteins.

**Data S3. (separate file)**

Comparison of gene set enrichment analysis (gene ontology biological process) in fresh (GSE210248) and cultured PASMC (GSE255669, GSE144274) and PAAF (GSE255669, GSE144932).

**Data S4. (separate file)**

List of differentially expressed genes in PASMC-PAAF co-culture model.
